# A dual role for the RNA helicase DHX34 in NMD and pre-mRNA splicing and its function in hematopoietic differentiation

**DOI:** 10.1101/2022.05.26.492072

**Authors:** Nele Hug, Stuart Aitken, Dasa Longman, Michaela Raab, Hannah Armes, Abigail R. Mann, Ana Rio-Machin, Jude Fitzgibbon, Kevin Rouault-Pierre, Javier F. Cáceres

**Affiliations:** MRC Human Genetics Unit, Institute of Genetics and Cancer, University of Edinburgh, Crewe Road South, Edinburgh EH4 2XU, UK; Centre for Genomics and Computational Biology, Barts Cancer Institute, Queen Mary University of London, London, United Kingdom; Centre for Haemato-Oncology, Barts Cancer Institute, Queen Mary University of London, London, United Kingdom

**Keywords:** DHX34, RNA helicase, NMD, pre-mRNA splicing, RNA targets, seCLIP, AML

## Abstract

The DExD/H-box RNA helicase DHX34 is a Nonsense-mediated decay (NMD) factor that together with core NMD factors co-regulates NMD targets in nematodes and in vertebrates. Here, we show that DHX34 is also associated with the human spliceosomal catalytic C complex. Mapping of DHX34 endogenous binding sites using Cross-Linking Immunoprecipitation (CLIP) revealed that DHX34 is preferentially associated with pre-mRNAs and locates at exon-intron boundaries. Accordingly, we observed that DHX34 regulates a large number of alternative splicing (AS) events in mammalian cells in culture, establishing a dual role for DHX34 in both NMD and pre-mRNA splicing. We previously showed that germline DHX34 mutations associated to familial Myelodysplasia (MDS)/Acute Myeloid Leukemia (AML) predisposition abrogate its activity in NMD. Interestingly, we observe now that DHX34 regulates the splicing of pre-mRNAs that have been linked to AML/MDS predisposition. This is consistent with silencing experiments in hematopoietic stem/progenitor cells (HSPCs) showing that loss of DHX34 results in differentiation blockade of both erythroid and myeloid lineages, which is a hallmark of AML development. Altogether, these data unveil new cellular functions of DHX34 and suggests that alterations in the levels and/or activity of DHX34 could contribute to human disease.

## INTRODUCTION

Nonsense-mediated decay (NMD) is an RNA quality control mechanism that targets mutated mRNAs harboring premature termination codons (PTCs) for degradation, but importantly also has a role in the regulation of cellular transcripts, in particular those associated with the stress response (Kurosaki et al. 2018; Karousis and Mühlemann 2019; Goetz and Wilkinson 2017). We previously identified *smgl-2 (smg lethal-2)*, an ortholog of human *DHX34* (DExH-box helicase 34), as a factor promoting NMD in *C. elegans* (Longman et al. 2007). We went on to show that this RNA helicase acts in the NMD pathway not only in nematodes, but also in zebrafish and in human cells and co-regulates NMD substrates with core NMD factors, such as UPF1 (Anastasaki et al. 2011; Longman et al. 2013). Mechanistically, DHX34 is recruited to the initial NMD surveillance complex via its interaction with hypo-phosphorylated UPF1. Subsequently, it promotes UPF1 phosphorylation, enhanced recruitment of UPF2 and dissociation of the ribosome release factor eRF3 from UPF1, which are all hallmarks of a transition to an NMD decay-inducing complex (Hug and Cáceres 2014).

Human DHX34 belongs to the DExH/D family of RNA helicases and harbors a helicase core formed by two (RecA)-like domains, a winged-helix domain (WH) and a helical bundle domain, known as the Ratchet domain (Sloan and Bohnsack 2018; Hug and Cáceres 2014). In addition, as with most DEAH box proteins, DHX34 also harbors a C-terminal OB (oligonucleotide/oligosaccharide binding fold) domain that can act to regulate conformational changes in the DEAH box helicases (Abdelhaleem et al. 2003; Ozgur et al. 2015; Hug and Cáceres 2014). A large majority of DExH/D proteins are RNA helicases that unwind RNA duplexes in an NTP-dependent manner and are involved in multiple aspects of RNA processing, including pre-mRNA splicing, ribosome biogenesis and mRNA translation (Jankowsky and Jankowsky 2000; Jankowsky and Bowers 2006). Furthermore, besides their role in RNA unwinding, they have been shown to remodel ribonucleoprotein complexes (RNPs) by removing proteins from RNA (Schwer 2001; Fairman et al. 2004; Jankowsky et al. 2001).

A common function for RNA helicases is in the process of pre-mRNA splicing, where eight conserved DExD/H RNA helicases have been shown to play essential roles in directing conformational rearrangements in the spliceosome. These include DDX46/Prp5, DDX39B/Sub2 and DDX23/Prp28 that belong to the DEAD-box family; DHX8/Prp22, DHX15/Prp43, DHX16/Prp2 and DHX38/Prp16 that belong to the DEAH-box family and SNRNP200/Brr2 that is part of the Ski-2 like family (Cordin and Beggs 2013; Bourgeois et al. 2016; De Bortoli et al. 2021). The function of these RNA helicases in constitutive splicing is diverse since they affect different steps of the spliceosomal cycle. The human spliceosome comprises five additional RNA helicases, which include SF3b125, DDX35, DDX41, eIF4AIII/DDX48 (a component of the Exon junction complex or EJC) and Aquarius (also known as intron-binding protein 160 or IBP60)(De et al. 2015). A role for several RNA helicases, such as DDX5 and DDX17, in alternative splicing has also been established (Hönig et al. 2002; Guil et al. 2003; Dardenne et al. 2014; Lee et al. 2018). The EJC fulfils a broader role in splicing regulation since it inhibits the use of cryptic splice sites, thus preventing the loss of exonic sequences (Boehm et al. 2018). Moreover, eIF4AIII/DDX48 affects the regulation of a large number of alternative exons (Michelle et al. 2012; Wang et al. 2014).

The fact that *smgl-2/DHX34* is essential for viability in nematodes, an organism where mutations in genes encoding core NMD factors are tolerated strongly suggested that SMGL-2/DHX34 fulfils at least one additional cellular function (Hug et al. 2016; Longman et al. 2007). Here, we show that DHX34, in addition to its established role in NMD, associates with the late spliceosome and impacts splicing regulation in mammalian cells in culture. We previously identified heterozygous germline variants in DHX34 in four families affected of inherited acute myeloid leukaemia (AML) and myelodysplastic syndrome (MDS) and showed that all these variants abrogated DHX34 NMD activity (Rio-Machin et al. 2020). Although *DXH34* is not mutated in sporadic AML, it is subject to alternative splicing in one third of sporadic cases, resulting in a premature stop codon that phenocopies germline mutations observed in familial patients with a broad impact on the AML transcriptome (Rivera et al. 2021). Due to the prevalence of mutations in spliceosomal proteins, such as DDX41, SF3B1, U2AF1 or SRSF2 in AML/MDS patients, it is tempting to speculate that DHX34 mutations and/or alternative splicing changes found in these patients could compromise not only its function in NMD, but also affect splicing events mediated by DHX34. Indeed, we show that DHX34 regulates AS of pre-mRNAs that have been linked to AML/MDS. Moreover, *DHX34* knock-down in hematopoietic stem/progenitor cells (HSPCs) demonstrated a disruption in erythroid and myeloid differentiation, potentially contributing to MDS/AML development.

In summary, we have unveiled a novel role for the RNA helicase DHX34 in alternative splicing regulation and showed that DHX34 is required for hematopoietic differentiation. These data highlight diverse cellular functions of DHX34 and suggest that alteration of its different RNA processing activities can contribute to human disease. These data highlight diverse cellular functions of DHX34 and suggest that alteration of its different RNA processing activities can contribute to human disease.

## RESULTS

### DHX34 interacts with complexes involved in mRNA processing

We previously showed that DHX34 binds directly to RNA and interacts with core NMD factors, including UPF1 and the Serine/Threonine-protein kinase SMG1, and also with proteins involved in other aspects of RNA degradation (Hug and Cáceres 2014; Melero et al. 2016). To investigate whether DHX34 is implicated in other steps of RNA biogenesis that extend beyond NMD and/or mRNA degradation, we sought to identify DHX34-interacting proteins. For this, we performed immunoprecipitation (IP) and mass spectrometry (MS) of anti-GFP DHX34 from a HEK293T cell line, where the endogenous locus had been tagged with a FLAG and GFP-tag using CRISPR/Cas9 genome editing (Fig. 1A). IP-MS profiles from three independent CRISPR clones, termed A5, A10 and 1B3, all displayed a significant enrichment for proteins involved in mRNA splicing, mRNA translation and Exon junction complex (EJC) components (Fig. 1B, C; Supplemental Table 1). Interacting proteins included the spliceosomal proteins PRPF19, ISY1, DDX41, the EJC components eIF4A3, MAGOH and ribosomal proteins RLA0 and RL4 (Fig. 1B-D). Importantly, all three independent clones exhibited a strong correlation of their interacting partners (Supplemental Fig. S1A). We also detected most of the DExD/H RNA helicases that components of the spliceosome in the DHX34 interactome, including DHX8/Prp22, DHX15/Prp43, DHX38/Prp16, and DDX41 (Fig. 1B-D; Supplemental Table 1). In agreement with our previous results, we confirmed the interaction of DHX34 with the NMD factor SMG1 and the no-go decay (NGD) factor Pelota (Harigaya and Parker 2010), as well as with ribosomal protein S6 (Supplemental Fig. S1B). Importantly, none of the DHX34-tagged clones significantly affected cell growth (Supplemental Fig. 1C).

**FIGURE 1.**
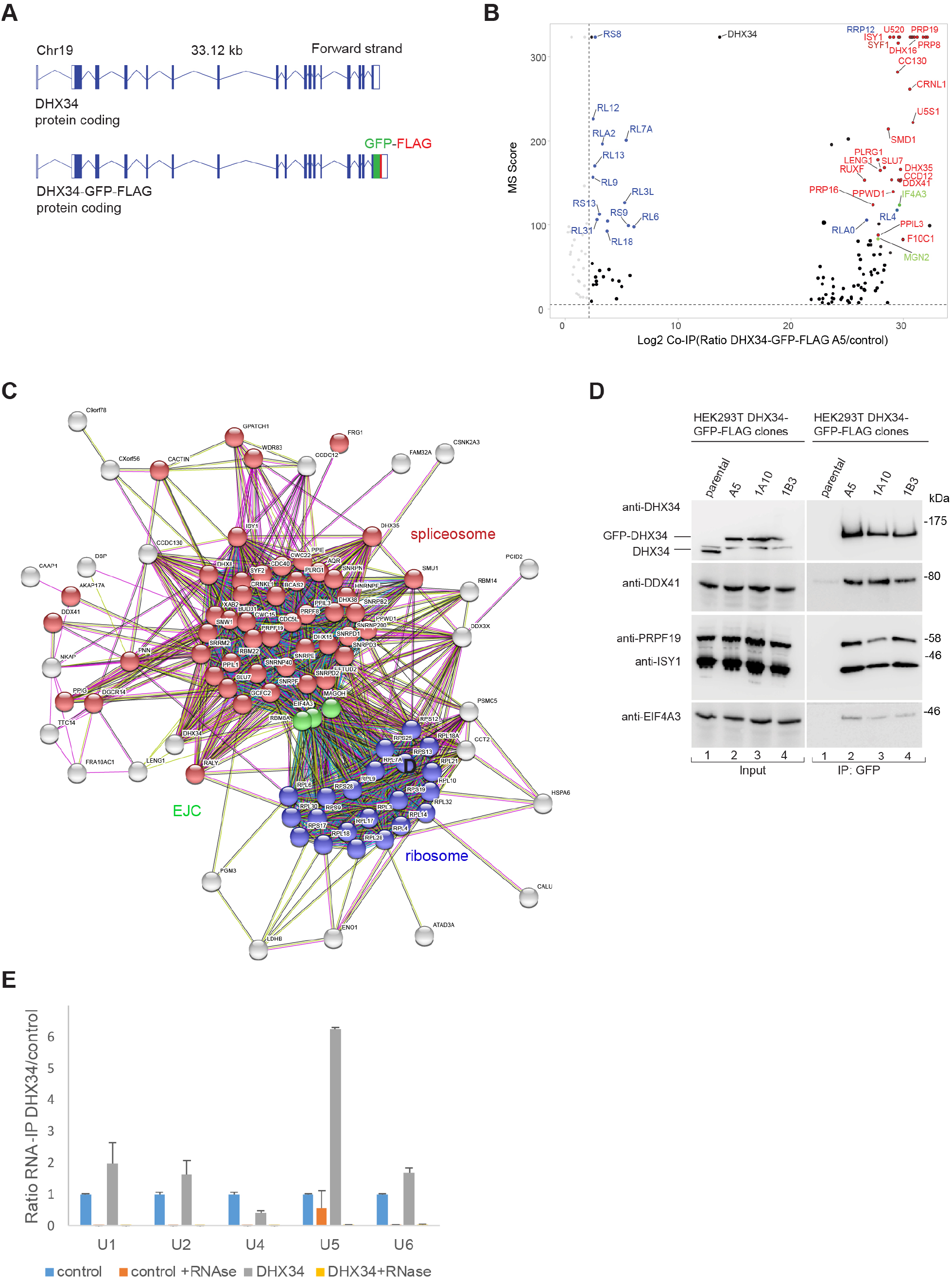
DHX34 interacts with mRNA processing complexes. (*A*) Cartoon depicting the CRISPR-mediated tagging of the endogenous DHX34 locus in HEK293T cells to generate C-terminal tagged DHX34-GFP-FLAG cell lines. (*B*) Volcano plot of 125 common interacting proteins identified by mass spectrometry (Log2 ratio >2) for DHX34-GFP-FLAG A5, 1A10 and 1B3 CRISPR clones. Protein names are indicated for the top 50 enriched ribosomal (blue), spliceosomal (red) and EJC proteins (green). Due to space constraints, not all protein names are indicated in the plot. All identified proteins are listed in Supplemental Table 1. (*C*) String network of interacting proteins identified by mass spectrometry of anti-GFP Immunopurifications from three independent CRISPR DHX34-GFP-FLAG clones. DHX34 interacts with protein complexes involved in mRNA biogenesis: spliceosome (red), EJC (green) and ribosome (blue). (*D*) Validation of mass spectrometry experiments with anti-GFP Immunoprecipitations (IPs) of three different CRISPR clones used for mass spectrometry. Inputs and anti-GFP IPs were separated by SDS-PAGE and probed with the indicated antibodies in Western blot assays. (*E*) U5 snRNA co-purifies with DHX34 whereas two other snRNAs present in the spliceosomal complex C, U2 and U6, are not enriched in the IP. RNA-protein complexes were immunopurified using anti-FLAG beads from GFP-FLAG A5 clone, following an elution step, RNA was reverse-transcribed and PCR amplified with specific primers for spliceosomal snRNAs.

The strongest enrichment of DHX34 interacting proteins was seen for proteins involved in the late spliceosomal reaction (complex C) (Fig. 1B-D; Supplemental Fig. S1D, E). Overall, 42 out of 49 annotated spliceoceosomal complex C proteins co-purified with DHX34 in the interactome, consistent with the finding that DHX34 was found to be dynamically associated with the spliceosomal complex C (Schmidt et al. 2014). As we previously showed that DHX34 is an RNA-binding protein (Hug and Cáceres 2014), we tested whether DHX34 interacts with spliceosomal small nuclear RNAs (snRNAs) by performing RNA Immunoprecipitation followed by RT-qPCR using the DHX34-GFP-FLAG clone A5 (Fig. 1E). Out of the five snRNAs tested, we only detected a strong enrichment of U5 snRNA in the RNA-Co-IPs, which disappeared upon RNase treatment (Fig. 1E). This is compatible with the observation that DHX34 interacts with the late spliceosome required for the second catalytic step where only U2, U5 and U6 snRNA are present in complex C and with the association of DHX34 with protein factors that are part of the U5 snRNP, such as PRPF8, SNRPD1/2 and 3, SNRNP200, SNRNP40, SNRNPE and SNRNPN (Fig. 1B, C). These findings indicate that DHX34 may influence various aspects of mRNA biogenesis and strongly suggest a role for DHX34 in pre-mRNA splicing.

### Genome-wide mapping of DHX34 binding sites using seCLIP

We have previously established that DHX34 is an RNA-binding protein using an mRNA capture assay (Hug and Cáceres 2014). In order to uncover the roles of DHX34 in pre-mRNA splicing and/or other aspects of RNA processing, we decided to focus on the identification of DHX34 RNA binding sites in the genome of the same cell line where the interactome was performed, HEK293T. DHX34 binding sites were identified using the seCLIP protocol (single-end enhanced crosslinking and immunoprecipitation) in the DHX34-GFP-FLAG A5 clone described above, using anti-GFP beads for the IP (Blue et al. 2022), with the parental cell line serving as a negative control. Purified RNA-DHX34 protein complexes were separated by SDS-PAGE (Fig. 2A) and cross-linked RNA fragments were shown to map predominantly to protein coding transcripts (Fig. 2B). Most non-protein coding transcripts identified with the seCLIP protocol were long non-coding RNAs (lincRNAs) and antisense RNAs (Supplemental Fig. S2A). Spliceosomal snRNAs were not detected and this most likely reflects the stringency of the RNase treatment during the seCLIP protocol. Using MEME (Bailey et al. 2009), we were unable to identify specific RNA-binding motifs (Supplemental Fig. S2D). This is in agreement with the poor sequence-specificity described for DExH/D RNA helicases that interact via their RecA domains with the sugar–phosphate backbone of RNAs (reviewed by (Bourgeois et al. 2016)). GO term analysis revealed DHX34 preferential binding to RNAs encoding splicing components (Supplemental Fig. S2E).

**FIGURE 2.**
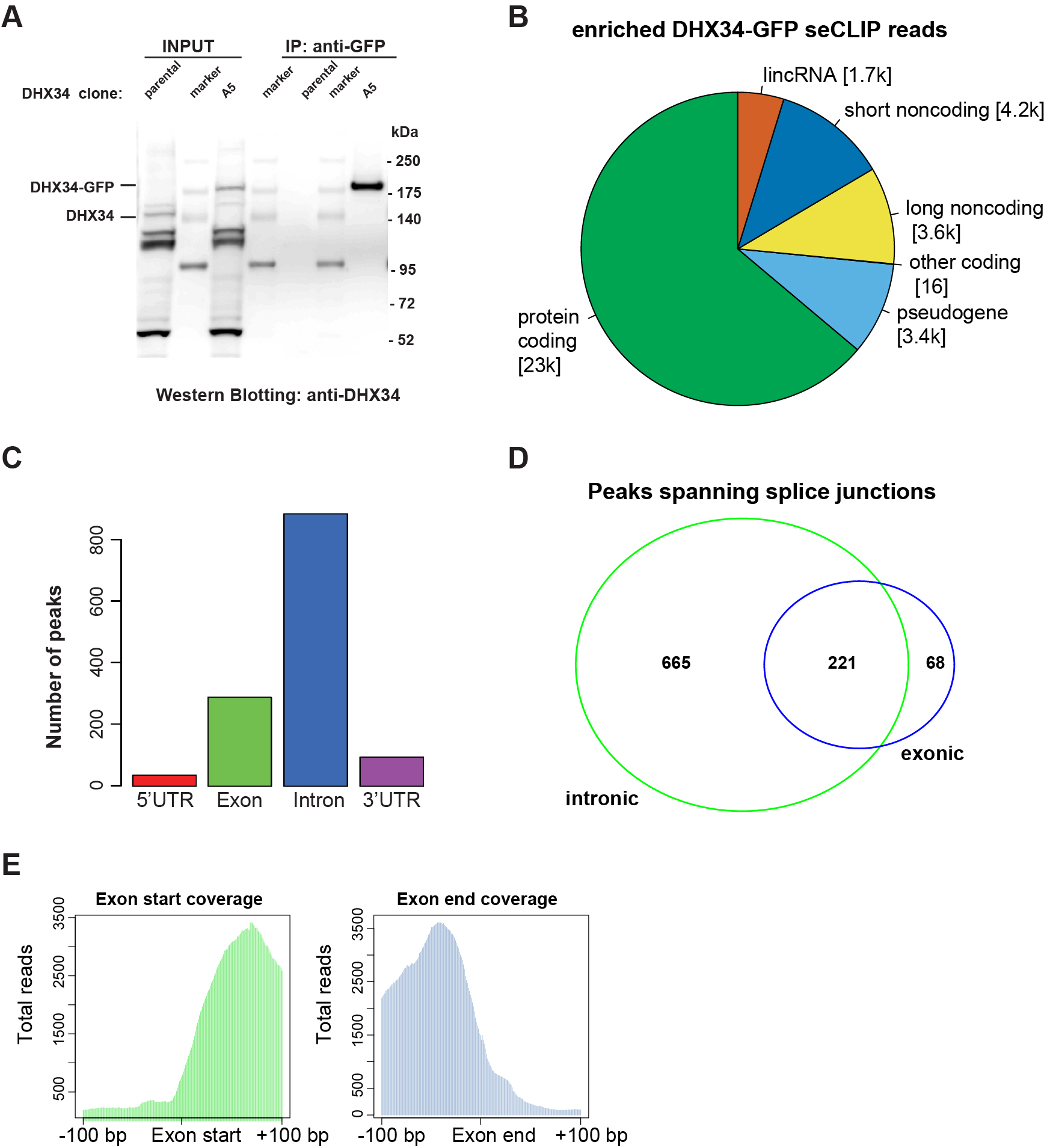
DHX34 binds in the proximity of exon-intron boundaries. (*A*) Western blot showing the samples used for the generation of the CLIP sequencing library. In each case, the band migrating at the size that corresponds to DHX34-GFP-FLAG was gel extracted and the RNA fragments were eluted. RNA sequencing libraries were constructed following the seCLIP protocol. Pie chart showing the distribution of DHX34 CLIP reads among different transcript species. Metagene analysis of DHX34 RNA binding. DHX34 binding peaks map mostly to introns and exons. Only few peaks were found in 5’UTR and 3’UTR. (*D*) Intronic and exonic peaks overlap suggesting that DHX34 binding spans splice junctions. (*E*) Pile up of DHX34 bound RNA reads at the exon-intron boundary.

We performed metagene analysis and found that DHX34 binding peaks mapped largely to exon and introns (Fig. 2C). For genes encoding at least three exons, DHX34 binding sites were mainly located in the mid-exons or introns, as expected. However, we observed higher-than expected DHX34 binding to first or last exon and first intron (Supplemental Fig. S2C). Crucially, a large number of peaks spanned splice junctions (Fig. 2D, E) strongly suggesting that DHX34 binds to pre-mRNA prior to co-transcriptional mRNA processing.

### Cellular pathways regulated by DHX34

To assess the global effects of DHX34 on the transcriptome of cells in culture, we performed RNA sequencing (RNA-seq) of HeLa cells that were depleted of DHX34 (Fig. 3A; Supplemental Fig. S3A).To extend our previous findings of DHX34 role in NMD (Hug and Cáceres 2014), we compared the upregulated transcripts upon DHX34 knock-down with those that were also upregulated upon depletion of the core NMD factor UPF1. We found that depletion of DHX34 affected the expression of 4,439 genes with 1,988 genes significantly overexpressed. Of these upregulated transcripts, 21% overlapped with previously identified UPF1 targets that were upregulated upon UPF1 knockdown (Longman et al. 2020)(Supplemental Fig. S3B). These upregulated targets that are co-regulated by DHX34 and UPF1 most likely represent bona fide NMD targets in HeLa cells (Fig. 3B; Supplemental Fig. S3B; Supplemental Table 2). Interestingly we noted that DHX34 depleted cells not only showed deregulation of cellular transcript levels, but also displayed changes in alternative splicing.

**FIGURE 3.**
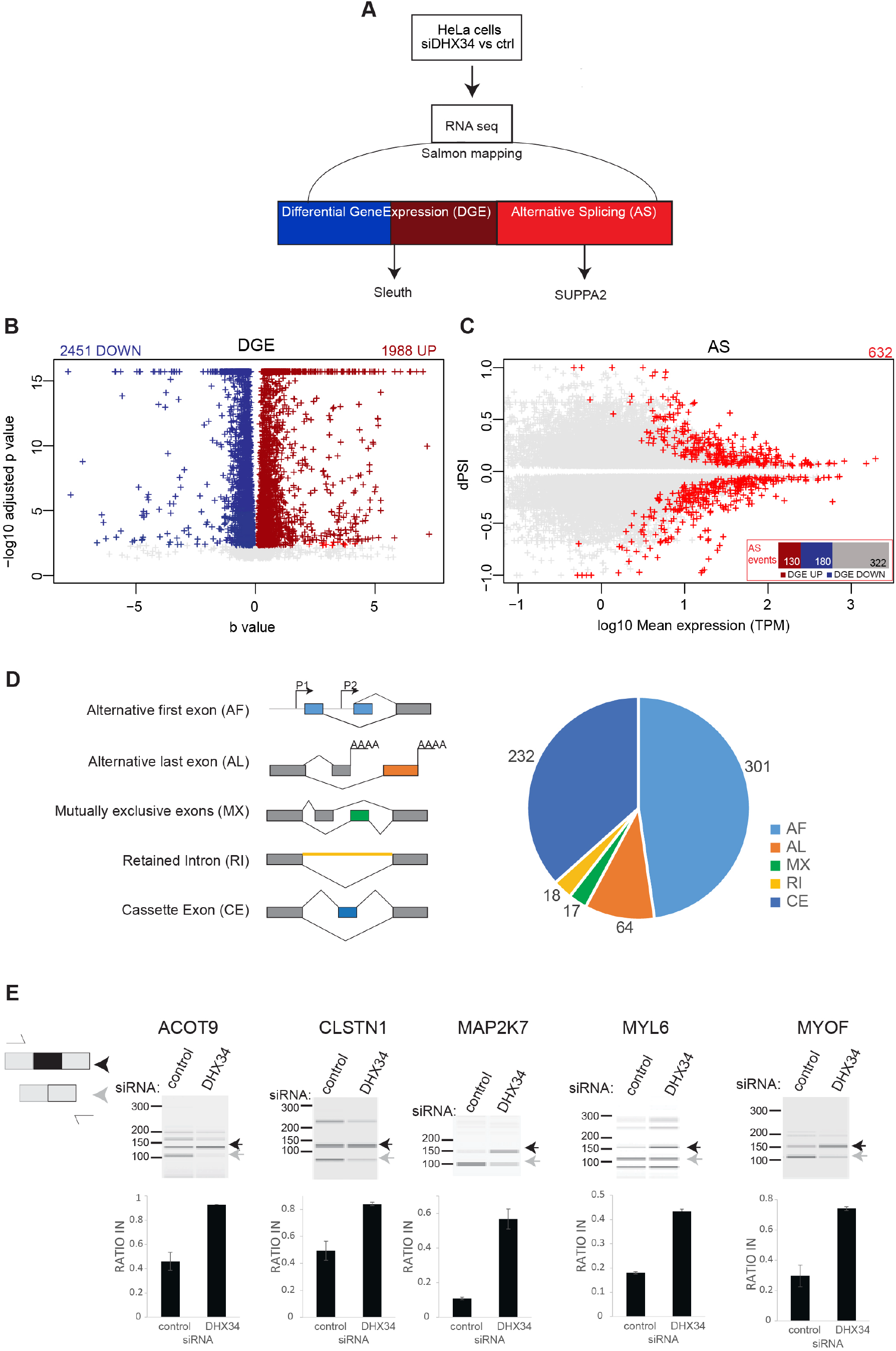
DHX34 regulates NMD and pre-mRNA splicing. (*A*) Outline of the experimental design for the analysis of changes in Gene expression and Alternative splicing upon DHX34 knock-down. RNA-seq was performed from HeLa cells depleted of DHX34 (siDHX34) or transfected with non-targeting siRNA pools (ctrl). Sequencing reads were mapped using Salmon and differential gene expression (DGE) was performed with Sleuth. Splicing changes were detected with SUPPA2. (*B*) Volcano plot of DGE changes upon DHX34 depletion are indicated by altered b-value and -log10 adjusted pvalue. (*C*) Splicing changes upon DHX34 depletion. Significant splice changes detected with SUPPA2 algorithm are depicted in red (dPSI>0.05, p≤0.05). Bar plot indicates pre-mRNAs that show AS changes as well as changes in gene expression: upregulated expression (DGE UP, dark red), downregulated (DGE DOWN, blue), not changed (grey). (*D*) Pie chart showing different types of alternative splicing events detected with SUPPA2. (*E*) Validation of cassette exon splice changes for ACOT9, CLSTN1, MAP2K7, MYL6 and MYOF transcripts by RT-PCR in HeLa cells depleted for DHX34 or treated with non-targeting siRNA (control). Dark grey arrow indicates transcript variant with included exon, light grey with excluded exon. Means from four individual data point obtained by RT-PCR using Bioanalyzer are plotted with standard deviations as error bars.

We measured “percentage spliced in” deltaPSI values using the SUPPA2 algorithm (Trincado et al. 2018), and detected 632 altered splicing events (Fig. 3C). Predominant changes were found in cassette exons (CE) (232 events) and alternative first exons (AF) (301 events) (Fig. 3D; Supplemental Table 2), with adj pvalue <0.05 and |deltaPSI|>0.05. To a much lesser extent, we also detected retained introns (RI), mutually exclusive exons (MX) and alternative last exons (AL) (Fig. 3D). We used RT-PRC analysis to validate AS regulation by DHX34 (CE) of 5 selected transcripts that displayed significant changes in the RNA-seq datasets. In all cases, DHX34 seems to promote skipping of the CE, since its knock-down leads to inclusion of alternative cassette exons in all tested pre-mRNAs (Fig. 3E). These results strongly suggest that DHX34 has dual role in HeLa cells, affecting both NMD and alternative splicing.

### Role of DHX34 in leukemia

We previously identified heterozygous mutations in *DHX34* in four families affected with inherited acute myeloid leukaemia (AML) and myelodysplastic syndrome (MDS) and showed that these mutations abrogated the NMD function of DHX34 using an NMD reporter (Rio-Machin et al. 2020). Interestingly, the *DHX34* pre-mRNA is subject to widespread alternative splicing in sporadic AML, which results in the inclusion of a poison exon harbouring a PTC, leading to a decrease in DHX34 mRNA levels due to alternative splicing coupled to NMD (AS-NMD)(Rivera et al. 2021). These findings strongly suggest that an altered activity of DHX34, by either mutation or AS-NMD, has a direct role in AML development. As a first attempt to investigate the functional role of DHX34 in blood disorders, we focused on the described role of DHX34 in NMD (Hug and Cáceres 2014) and in pre-mRNA splicing (this study) in a more relevant cellular system. For this, we performed RNA sequencing (RNA-seq) of the immortalized K562 myeloid leukaemia cell line following depletion of *DHX34*, which was verified by qRT-PCR (Supplemental Fig. S4A, B) and changes in gene expression (DGE) and in alternative splicing were assessed (Fig. 4A-C; Supplemental Fig. S4C, D).

**FIGURE 4.**
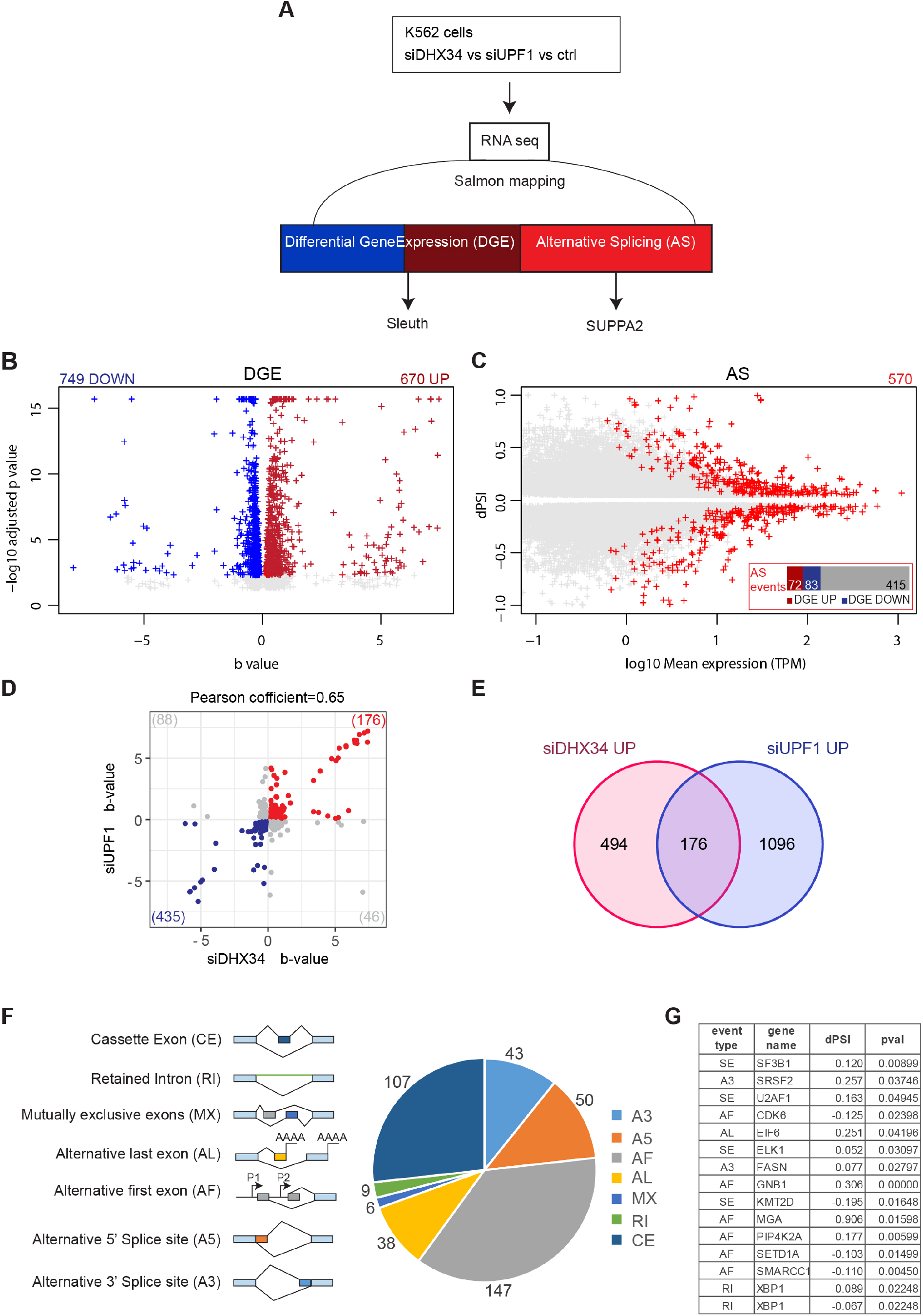
DHX34 is an NMD factor and regulates pre-mRNA splicing in K562 cells. *(A)* Outline of experimental design for RNA sequencing and analysis. RNA seq was performed for DHX34 knockdown (siDHX34), UPF1 knockdown (siUPF1) and compared to a non-targeting siRNA (ctrl). Sequencing reads were mapped using Salmon and differential gene expression (DGE) was performed with Sleuth. Splicing changes were detected with SUPPA2. *(B)* Volcano plot of DGE changes upon DHX34 depletion are indicated by altered b-value and -log10 adjusted pvalue. *(C)* Splicing changes upon DHX34 depletion. Significant splice changes detected with SUPPA2 algorithm are depicted in red (dPSI>0.05, p≤0.05). Bar plot indicates pre-mRNAs that show AS changes as well as changes in gene expression: upregulated expression (DGE UP, dark red), downregulated (DGE DOWN, blue), not changed (grey). *(D)* Scatter plot of the correlation between expression changes in DHX34 and UPF1 depletion. Each dot represents a common differentially expressed gene. Genes significantly upregulated in both DHX34 and UPF1 are labeled in red, genes which are downregulated in blue. *(E)* Venn diagram showing the number of common transcripts up-regulated (UP) in DHX34 and UPF1 knockdown cells. *(F)* Pie chart showing different splicing events upon DHX34 knockdown detected with SUPPA2. (*G)* Table listing AS events in genes linked to AML.

First, we focused on the role of DHX34 in NMD and compared the effects of depleting the core NMD factor UPF1with DHX34 depletion. We had previously used microarray profiling to show that DHX34 and UPF1 co-regulate a significant group of mRNA transcripts in nematodes, zebrafish and HeLa cells (Longman et al. 2013; Hug and Cáceres 2014).

Importantly, we validated these previous observations in K562 cells, with 26% of RNAs upregulated upon DHX34 depletion (176/670), being also upregulated upon knock-down of UPF1 (Fig. 4D, E; Supplemental Table 3), displaying a robust co-regulation (Pearson’s correlation r=0.65, p<0.0001) (Fig. 4D). These results clearly show that DHX34 is a general regulator of NMD in K562 cells and provide a list of potential NMD targets for this RNA helicase.

Interestingly, as observed with HeLa cells, DHX34-depleted K562 cells also displayed changes in alternative splicing (Fig. 4C, F). We measured “percentage spliced in” deltaPSI values using the SUPPA2 algorithm (Trincado et al. 2018), and detected 570 splicing changes (Fig. 4C, F; Supplemental Table 3). The most predominant changes were in alternative first exons (AF) (147 events) and in cassette exons (CE) (107 events) (Fig. 4F) with adj pvalue <0.05 and |deltaPSI|>0.05. Importantly, depletion of DHX34 led to differential splicing of several pre-mRNAs in genes that are are frequently mutated in MDS/AML, including SF3B1, SRSF2 and U2AF1 Fig. 4G).

### DHX34 regulates its own pre-mRNA splicing

It was recently shown that DHX34 is subject to widespread alternative splicing in sporadic AML, resulting in the inclusion of alternative exon 12b that harbors a PTC, leading to Alternative splicing coupled to NMD (AS-NMD) (Rivera et al. 2021)(Fig. 5A). Since we unveiled a dual role for DHX34 in NMD and AS regulation, we decided to explore whether DHX34 exerts a regulation of of its own pre-mRNA splicing.

**FIGURE 5.**
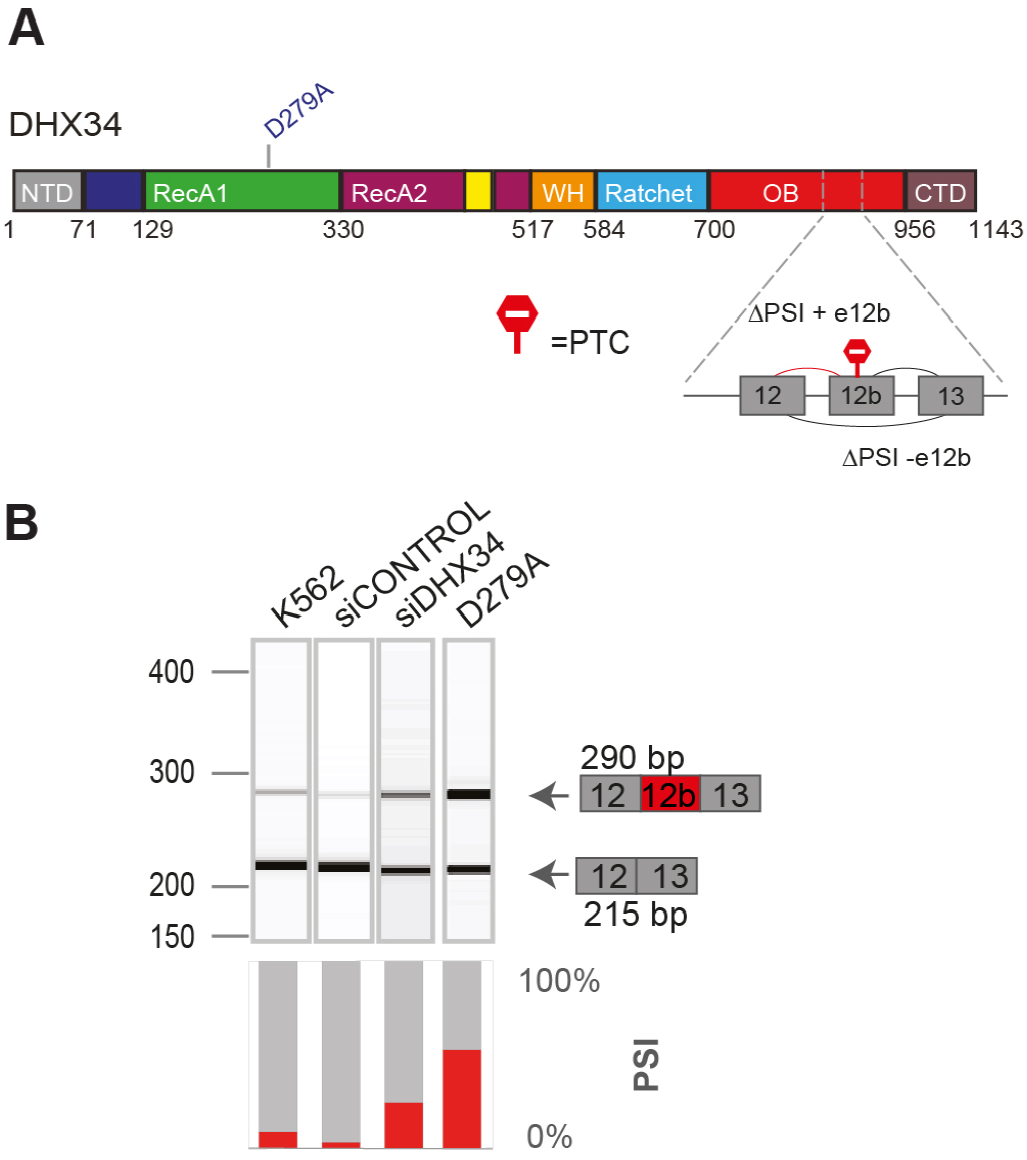
DHX34 regulates pre-mRNA splicing of its own pre-mRNA in K562 cells. *(A)* Schematic of DHX34 protein domain structure, including the D279A catalytic mutation. Part of the *DHX34* pre-mRNA exon-intron structure, including the alternative exon 12b harboring a PTC, is depicted. *(B)* RT-PCR analysis of splicing patterns of DHX34 exon 12b in knockdown and mutant conditions. RT-PCR products were resolved using Bioanalyzer (top panel) and relative splicing changes (PSI) were quantified.

RT-PCR analysis of endogenous pre-mRNA in K562 cells upon siRNA-mediated knock-down of DHX34 revealed an increase in the isoform containing E12b (Fig. 5B; Supplemental Fig. S5). Interestingly, this result was confirmed in an engineered K562 catalytic mutant cell line, where a mutation was introduced in an aspartate (D) residue in the Walker B motif (p.D279A in Motif II) that is required for ATP hydrolysis (Hanson and Whiteheart 2005; Hug and Cáceres 2014). In addition, RNA-seq analysis of K562 cells upon depletion of *DHX34* or harboring the p.D279A catalytic mutation confirmed these findings (Supplemental Fig. S5). Altogether, these results unveil a role for DHX34 in the regulation of its own expression and suggest the existence of an elaborate feed-back mechanism by which DHX34 could prevent the expression of the isoform containing exon 12b via NMD and/or alternative splicing, maintaining appropriate levels of DHX34 protein.

### A role for DHX34 in hematopoiesis

Finally, to gain further insight into the role of DHX34 in hematopoiesis, we used a lentiviral approach to generate a knock-down of *DHX34* in hematopoietic stem/progenitor cells (HSPCs) isolated from human umbilical cord blood (Supplemental Fig. S6). A significant knock-down was observed in transduced CD34^+^ cells (Supplemental Fig. S6B). Cells were sorted by flow cytometer (Dapi^-^CD34^+^GFP^+^) and placed in expansion medium where they showed a lower proliferation rate at day 7 (Supplemental Fig. S6C). Next, sorted cells were grown in semi-solid medium to assess the capacity of progenitors to proliferate and differentiate into the different myeloid and erythroid lineages/colonies. Interestingly, *DHX34* knock-down cells demonstrated an impaired capacity to generate colonies in both erythroid lineage (Burst Forming Units: BFU-E) and myeloid lineage (Colony Forming Units granulocytic-granulo/monocytic-monocytic: CFU-G/GM/M) (Fig. 6A), while no significant apoptosis was detected (Supplemental Fig. S6D).

**FIGURE 6.**
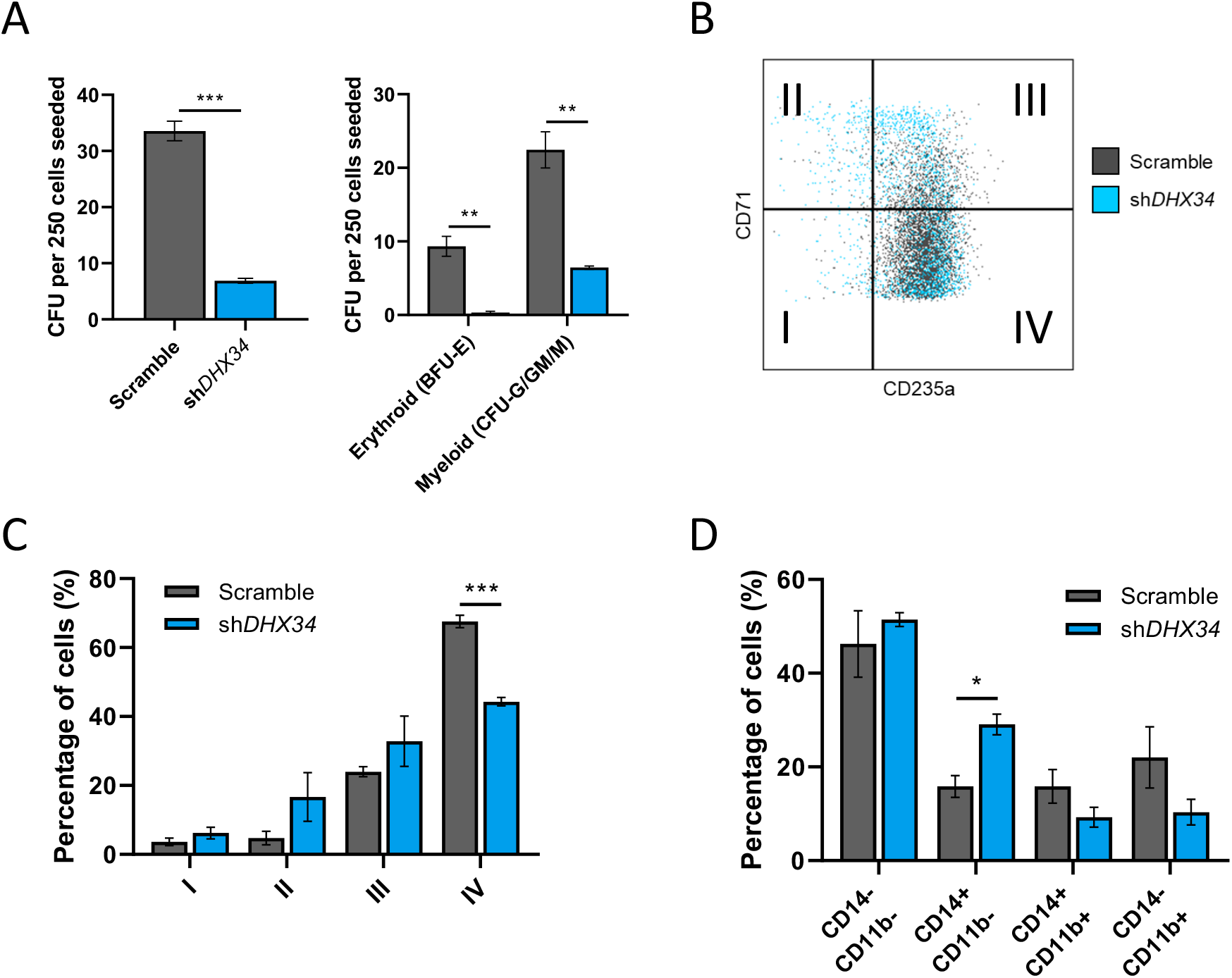
Loss of DHX34 impairs HSPC differentiation. (*A*) Cells were sorted by FACS (Dapi^-^ CD34^+^GFP^+^) and placed in methylcellulose prior to being scored at day 14. Left panel represents total colonies scored and right panel colonies scored in erythroid (BFU-E) and myeloid (CFU-G/GM/M) lineages (n=3). (*B*) FACS plot representing cells at day 14 in erythroid conditions, blue represents cells KD for *DHX34*, and grey represents control cells (scramble). (*C*) Bar chart quantifying the percentage of erythroid differentiated cells in the different quadrants (I, II, III & IV displayed on panel (B)). Cells were sorted by flow cytometer (Dapi^-^CD34^+^GFP^+^), cultured in erythroid differentiation conditions and immunophenotyped by FACS at day 14 based on CD71 and CD235a expression (n=3). (*D*) Bar charts representing the percentage of granulo-monocytic cells after Dapi^-^CD34^+^GFP^+^ cells were sorted by flow cytometer and cultured in granulo-monocytic conditions for two weeks. At day 14 cells were immunophenotyped by FACS based on CD11b and CD14 expression (n=3). *=p<0.05; **=p<0.01; ***=p<0.005

In light of these phenotypes, we investigated the impact of the loss of expression of *DHX34* during erythropoiesis and granulo-monocytic differentiation. Dapi^-^CD34^+^GFP^+^ were cultured under erythroid conditions and immunophenotyped at day 14 with CD71 (transferrin receptor) a marker of early erythroid differentiation and CD235a (Glycophorin A) a marker of mature erythroid cells.

Strikingly the knock-down cells demonstrated a significant blockage in erythroid terminal differentiation (Fig. 6B, C). When cells were placed in granulo/monocytic conditions, *DHX34* depleted cells showed an increase in CD14 expression (Fig. 6D), which is usually expressed by blast AML cells.

These findings reveal that DHX34 downregulation leads to ineffective erythropoiesis, which is a hallmark of AML and increased expression of CD14, which is often seen at the surface of AML blasts.

## DISCUSSION

DExH/D RNA helicases are involved in almost every aspect of RNA processing from RNA synthesis in the nucleus until mRNA translation and degradation in the cytoplasm. It is also common that individual helicases could be involved in more than one aspect of RNA processing, such as DHX9, which has been linked to alternative splicing, RNA export and miRNA biogenesis and function (reviewed by (Bourgeois et al. 2016)). We previously established a mechanistic role for DHX34 in the NMD pathway by showing that this RNA helicase promotes the transition from the initial NMD complex that surveys the presence of a PTC (SURF complex) to a Decay-inducing complex (DECID) where the actual RNA degradation occurs (Melero et al. 2016; Longman et al. 2013; Hug and Cáceres 2014). In this study, we identified an additional role for DHX34 in splicing regulation. We confirm DHX34 as a component of the catalytic spliceosomal complex C and show that DHX34 predominantly binds to pre-RNA in the vicinity of intron-exon junctions and has a role in the regulation of alternative splicing (Figs. 2-4). Interestingly, DHX34 was identified as a candidate neurodevelopmental gene; raising the possibility that this could be linked to its function in NMD, pre-mRNA splicing or another yet to be identified cellular function (Paine et al. 2019). The NMD and AS functions of DHX34 could operate independently; however, we show here that a subset of pre-RNAs that undergo AS changes upon DHX34 knock-down are also upregulated, and in K562 cells show co-regulation by UPF1 (Figs. 3 and 4).

The spliceosome undergoes extensive conformational and compositional rearrangements that are catalyzed by eight RNA helicases of the DExD/H family (De Bortoli et al. 2021). The activity of DEAH RNA helicases, such as DHX34, is often regulated through G-patch proteins, which function as adaptors that recruit them to functional sites and enhance their activity (Studer et al. 2020). The G-patch domain is an intrinsically unstructured region containing a set of conserved glycines that interact with an auxiliary OB-fold (oligonucleotide/oligosaccharide-binding fold) of their cognate DEAH box helicase and mediate protein-protein and RNA-protein interactions (Bohnsack et al. 2021; Robert-paganin et al. 2015; Studer et al. 2020). The interactome of DHX34 in HEK293T cells revealed the presence of one such protein, GPATCH1 (Fig. 1A). A recent study revealed that GPATCH1 copurifies with DHX35 and with components of catalytically active spliceosomes, strongly suggesting that the GPATCH1/DHX35 pair functions together to promote splicing fidelity (Sales-Lee et al. 2021). Whether GPATCH1 also acts together with DHX34 in pre-mRNA splicing; or and/or whether DHX34 is regulated by a different GPATCH protein, remains to be determined.

Previously, we also showed that DHX34 interacts with of RUVBL1-RUVBL2 AAA-ATPases and regulates their activity by stabilizing a conformation that does allow nucleotide binding and thereby down-regulates ATP hydrolysis of the complex (López-Perrote et al. 2020). Interestingly, RUVBL1 and RUVBL2 are essential constituents of several additional large complexes, with functions in chromatin remodeling. They are also part of the R2TP complex, a HSP90 co-chaperone involved in the assembly and maturation of large complexes that include RNA polymerase II, the Phosphatidylinositol 3-kinase-related kinase (PIKK) family members and the spliceosome (Dauden et al. 2021). The HSP90/R2TP complex, together with the ZNHIT2 cofactor, has a role in binding unassembled U5 proteins, including PRPF8, EFTUD2 and SNRNP200 in the cytoplasm and promotes the formation of the U5 snRNP particle (Malinová et al. 2017). Thus, it would seem plausible that DHX34 may have a role via the R2TP complex in regulating U5 snRNP. Indeed, we have previously shown that DHX34 does interact with PRPF8 (Hug and Cáceres 2014) and binds to U5 snRNA (Fig. 1D). Future work will determine whether DHX34 regulation of RUVBL1/2 has an impact on the R2TP complex and on U5 snRNP function.

A role for DHX34 in inherited acute myeloid leukaemia (AML) and myelodysplastic syndrome (MDS) was initially suggested by the presence of heterozygous mutations in *DHX34* in four families affected with this blood disorder, which affected its role in NMD (Rio-Machin et al. 2020), It was recently shown that another way of inactivating DHX34, beyond mutations, is by changes in alternative splicing coupled to NMD (AS-NMD) (Rivera et al. 2021). We show here that DHX34 downregulation leads to ineffective erythropoiesis, which is a hallmark of AML (Fig. 6). The discovery that DHX34 is involved in the regulation of pre-mRNA splicing is suggestive in regards to its function in AML/MDS, since recurrent mutations found in myeloid malignancies including genes encoding splicing factors, LUC7L2, RBM39, SF3B1, SRSF2, and U2AF1(Rahman et al. 2020; Zhang et al. 2015; Lee et al. 2016; Inoue et al. 2016; Wang et al. 2019; Daniels et al. 2021). This could be a common theme in blood disorders since deregulation of splicing factors was also found in pediatric B-cell acute lymphoblastic leukemias (B-ALL) (Black et al. 2018). In the case of DHX34, we observed predominantly effects on the regulation of cassette exons (CE), as well as the selection of first exons (AF), likely to impact on the choice of promoters (Figs. 3D, 4F). Mutations in another RNA helicase, the DEAD-box protein DDX41, have been identified in familial and acquired cases of myelodysplasia and acute myeloid leukemia, and these mutations in *DDX41* also give rise to defects in pre-mRNA splicing (Polprasert et al. 2015). Alterations in DHX15, another DExD/H-box RNA helicase that is part of the spliceosome and also functions in the ribosome biogenesis were also identified in an AML cohort (Faber et al. 2016). The recent finding that DHX34 is subject to alternative splicing in sporadic AML leading to the inclusion of a poison exon that results in AS-NMD highlighted the fact that DHX34 can be inactivated in familial AML not only via mutation but also through alternative splicing regulation (Rivera et al. 2021). Interestingly, we show that DHX34 regulates the abundance of its own pre-mRNA, via a mechanism involving AS-NMD that consequently reduces levels of DHX34 protein, this highlighting its role in maintaining cellular homeostasis (Fig. 6)

In summary, we have uncovered a dual role for the DExH/D-box RNA helicase, DHX34, in NMD and in the regulation of pre-mRNA splicing. Importantly, we show that DHX34 is required for the proper differentiation of HSCs to the erythroid lineage and myeloid lineage, which can possibly be explained by its role in NMD and/or in AS regulation.

## MATERIALS AND METHODS

### Cell Culture and transfections

HeLa and HEK293T cells were maintained in DMEM media with high glucose, GlutaMAX™ Supplement, pyruvate (Gibco Life technologies; 10569010) supplemented with 10% FCS, 1% penicillin/streptomycin at 37°C in the presence of 5% CO_2_. DHX34-FLAG-GFP clones were maintained in the same media. Cells were grown without antibiotic prior to transfections, which were carried out in Opti-MEM reduced serum medium (Gibco, 31985047). K562 cells were maintained in RPMI (GIBCO) media supplemented with 5mM Glutamin, 10% FCS and 1% penicillin/streptomycin at 37°C in the presence of 5% CO_2_. Transfections of siRNA oligos were done using DharmaFECT 1 (Dharmacon, T-2001-03) following manufacturer’s protocol (Supplemental Table 4). For total RNA-sequencing cells were plated in 6-well plates and transfected with 50 pmol of indicated siRNAs. Cells were expanded into 10 cm plates the following day and were transfected with 150 pmol of the same siRNAs on day 3 and were harvested for analysis 4 days after the first depletion.

### Design and screening of CRISPR cell lines

guideRNAs (gRNAs) were designed using sgRNA Designer CRISPRko (Broad institute, https://portals.broadinstitute.org/gpp/public/analysis-tools/sgrna-design) and CHOPCHOP https://chopchop.cbu.uib.no/Cas-Designer. Guides were cloned into pSpCas9(BB)-2A-Puro (px459) V2.0 (Ran et al., 2013). For GFP-FLAG tagging the repair template containing synthetic homology arms, 3XFLAG-tag and eGFP (amplified from phrN1GFP) was cloned into the pCDNA3.1(+) backbone using Gibson assembly. For introduction of point mutations into DHX34 by HR ssDNA oligos were used with mutated PAM sites. The gRNA/Cas9 plasmid and linearized repair template were transfected and selected with 1 1g/ml puromycin for 48 hours. 5 days post-transfection surviving cells were cloned into 96 well plates and expanded. Colonies were PCR screened and correct targeting verified by Sanger sequencing. For base editing gRNAs were cloned into pSPgRNA and transfected together with AncBE4max-P2A-GFP (Koblan et al. 2018)(ratio 1:3). Five days post-transfection GFP-positive cells were sorted by FACS into 96 well plates. Target regions were PCR amplified and base editing verified by Sanger sequencing. Sequences for templates and sgRNAs are listed in Supplemental Table 4.

### Mass Spectrometry

Cells were harvested and lysed as in immunoprecipitation protocol (see below). α-GFP antibody-coupled magnetic beads (Chromotek) were equilibrated with IP buffer. Lysates were resuspended in 500 μl IP buffer for capture of DHX34-FLAG-GFP bound proteins and subsequent mass spectrometry analysis. Immunoprecipitation were performed on Kingfisher Duo robot (Thermo) and subjected to in solution digestion according to standard protocols. for 4 h. Fractionated peptides were separated and analyzed using a Dionex RSLC Nano system coupled to a Thermo Q-Exactive Plus instrument. (Thermo Fisher Scientific). Raw MS data were analyzed using MaxQuant (v 1.5.6.5) (Max Planck Institute of Biochemistry) in conjunction with UniProt human reference proteome release 2016\_11 (uniprot.com), with match between runs (MS/MS not required), LFQ with 1 peptide required, and statistical analyses performed in R (RStudio 1.1.453 / R x64 3.4.4) (rstudio.com) using Wasim Aftab’s LIMMA Pipeline Proteomics (github.com/wasimaftab/LIMMA-pipeline-proteomics) implementing a Bayes-moderated method. Interactome analysis including gene ontology was carried out by inputting protein list into STRING (string-db.org/) and Gene Ontology enRIchment anaLysis and visuaLizAtion (GOrilla) (http://cbl-gorilla.cs.technion.ac.il/) (Eden et al. 2009).

### Immunoprecipitation and Western Blotting

Cells were washed and harvested in ice-cold PBS before pellets were lysed with immunoprecipitation (IP) buffer (20 mM Tris-HCl pH 8, 150 mM NaCl, 1mM EDTA, 1% NP-40, 0.2% Deoxycholate, Complete Protease Inhibitor (Roche), Phospho STOP (Roche), 1 mM DTT) for 20 min on ice. Cell lysates were treated with 80 1g/ml RNase A per 1 ml of extract. Anti-GFP MA (Chromotek) magnetic beads were washed, bound proteins were eluted with NuPAGE LDS sample buffer supplemented with reducing agent (Thermofisher). Proteins were resolved by SDS-PAGE on NuPAGE 3-8% Tris-Acetate precast gels (Thermofisher) and protein transfer was achieved using the iBlot™ 2 Gel Horizontal Transfer Device (Thermofisher). Nitrocellulose membranes were blocked in 5% BSA in PBS/Tween 20 (0.1%) and probed with the appropriate primary antibody diluted in blocking solution 1:1000. HRP-conjugated secondary antibodies (BioRad) were used at 1:10,000 and blots developed with ChemiGlow detection reagent (ProteinSimple) and visualized using ImageQuant LAS 4000 chemiluminescent camera (GE Healthcare). For RNA-Immunoprecipitations FLAG-Immunoprecipitation from DHX34-GFP-FLAG A5 cells were performed with in NET2 buffer (50 mM Tris-HCl pH7.5, 150mM NaCl, 0.05% Triton-X-100) using M2 agarose. Half of the samples were treated with 80 1g/ml RNase A as a negative control. IPs were washed 8X with NET2 and bound proteins eluted with 3XFLAG peptides and extracted with Trizol (Thermofisher). Eluted RNA was reverse transcribed and amplified with spliceosomal RNA specific primers (Supplemental Table 4)

### Antibodies

Anti-DDX41 (15076, Cell Signaling), Anti-PRP19 (ab27692, Abcam), Anti-ISY1 (HPA016995, Atlas Antibodies), Anti-SMG1 (ab30916, Abcam), Anti-UPF1 (# A300-036A, Bethyl), Anti-Pelota (bs-7821R, BioSS) Anti-RP6 (2217, Cell Signaling), Anti-GFP (ab290, Abcam), Anti-Tubulin (# 4026, Sigma-Aldrich), Anti-DHX34 is a peptide-specific antibody raised against human DHX34 obtained from Eurogentec (Hug and Cáceres 2014). For Immunopurifications GFP-Trap-MA beads (Chromotek) and Anti-FLAG affinity gel (A2220, Sigma) were used.

### seCLIP protocol

seCLIP experiments were performed following a published protocol (Blue et al. 2022), with minor modifications. The immunoprecipitation (IP) step was carried out using an anti-GFP beads (Chromotek). Five independent experiments for DHX34 were performed using DHX34-GFP-FLAG A5 clone with parental HEK293T serving as negative control. The five DHX34 seCLIP libraries and negative controls with different barcodes were pooled together and sequenced on a single lane by single end sequencing 50nt together with on an Illumina *HiSeq* 2000 system (Wellcome Trust Clinical Research facility at the University of Edinburgh (WTCRF)). Equivalent Input the control libraries were sequenced on a different lane. The seClip bioinformatics protocol was followed with adaptations to account for the adaptors and sequencing technology used here (Blue et al. 2022). Briefly, fastq files were merged then 3’ adaptors (starting with InvRand3Tr3) were trimmed using cutadapt (Martin, 2011), then the 5’ adaptors trimmed from reads lacking the 3’ adaptor (starting with InvAR17) and Illumina adaptors trimmed from the remaining reads. UMIs were identified in all three sets of reads using umi_tools (Smith et al. 2017). Read sequences were reverse-complemented prior to, and following, trimming and UMI processing using seqkit (Shen et al. 2016) to account for the forwards-reverse orientation of the reads. Reads were aligned to the human genome (GRCh38 93) using bowtie2 (Langmead and Salzberg 2012), sorted and indexed using samtools (Li et al. 2009), deduplicated using umi_tools and again sorted and indexed. The three sets were merged, and uniquely mapping reads retained, reads mapping to transposable elements were removed. The mapping rate was determined at all steps of processing. The final number of mapped reads for DHX34 GFP replicates ranged from 106k to 537k (0.4% to 9.3% of total reads over the replicates) totalling 6,926k reads. Of DHX34 GFP reads, 75% had either the 3’ or 5’ adaptor, 26.5% of all DHX34 GFP reads were retained after the removal of duplicates, reducing to 4.7% after the removal of multiply-mapping reads and those mapping to repeats (seCLIP_mapping.xlsx). To obtain an overview of the mapping of DHX34, the distribution of GPF IP reads across biotypes was assessed. Total normalized reads mapping to gene bodies per biotype is shown in pie_chart_biotypes_GFP_IP.pdf for all GFP IP reads, and when summing reads over those genes where GFP IP is 1.5 times enriched over negative GFP IP, enriched over the input and over both. With and without filtering for enrichment, DHX34 predominantly maps to protein-coding genes. In these charts, the specific short non-coding biotypes (snoRNA, snRNA etc) are aggregated under short noncoding, and similarly for pseudogenes and long noncoding other than lincRNA which is shown. Peaks were called in DHX34 GFP samples using macs2 (Zhang et al. 2008), with the merged negative GFP inputs as control (options: --broad-cutoff 0.1 -g hs -- nomodel --extsize 100) for each replicate individually and for merged DHX34 GFP replicates. Peaks with -log10 p value of at least 10 were retained, and a bed file of the union of peaks from all replicates was created. To review the absolute raw read count support for the peaks, the number of reads mapping to these peaks, and to the surrounding region (+/-the peak width) was quantified using htseq-count (Anders et al. 2015). The peak count to region count ratio was used to filter out regions of non-specific mapping. 1084 peaks with at least 5 reads in the merged data, and where the ratio of peak count to region count was greater than or equal to the mean (0.89) were selected for further consideration. (seClip_peak_to_region_depth.pdf). Peaks were reviewed manually from snapshots created from the IGV browser. Of the peaks selected, 957 (88%) had 5 or more reads in one or more individual replicates in addition to the calling of the peak in the merged data. 238 (22%) had this support in two or more replicates. The correlation of raw counts for the selected peaks across replicates was fair (seClip_replicate_correlation.pdf Supplemental Fig. S2B), with the exception of GFP_IP1. Peaks in GFP_IP1 though strongly indicated, were less well replicated. The selected peaks were supported by 5-11 raw reads across replicates (1^st^-3^rd^ quartile) raw counts (truncated at 11 to suppress outlying counts) are shown in the heatmap seClip_replicate_heatmap_0_11.pdf.SFig 3D Peak widths were 102-108 bases (1^st^-3^rd^ quartile).

### Gene expression profiling: RNA extraction, library preparation and RNA-sequencing

Total RNA was isolated from depleted cells and CRISPR clones using RNAeasy kit (Qiagen) and treated with TURBO DNA-free™ DNase I kit (Invitrogen Ambion; AM1907). Libraries were prepared by BGI (Hela cells RNA samples) or Novogene (K562 siRNA treated and CRISPR edited clonal RNA samples).

### RNA-sequencing analysis

Transcript abundances were quantified using salmon (Patro et al. 2017) from a transcriptome index compiled from coding and non-coding cDNA sequences defined in Ensembl GRCh38 93 (salmon version 1.5.2; using the flags --gcBias --numBootstraps 100). Differential expression was called with the sleuth R package (Pimentel et al. 2017) (significance taken as q<=0.005). Additional analyses to assess the consistency of the direction of expression change were performed using the Wald test in sleuth (significance taken as q<=0.005). Each condition (3 biological replicates per clone) was compared to wild type K562 (or SCR in the case of DHX34 and UPF1). PCA plots were generated for each comparison, and for all K562 and both CRISPR datasets combined. Genes consistently upregulated or consistently downregulated (significant at the gene level and all significant transcript level changes in the same direction) were taken forward for further analysis. Annotated splicing event occurrence (including alternative 3’ and 5’ splice site usage, exon skipping and alternative first/last exon usage) was assessed by Suppa2 version 2.3 (Trincado et al. 2018). Splicing analysis used the same transcript models and gene annotation as for differential expression calling. R scripts were written to filter, format and integrate the results. The default level of statistical significance of q<=0.05 was used in SUPPA2 analyses. A dPSI of >=0.05 and isoform expression of >=0.5 TPM were required in addition in SUPPA2 calls. LSVs were considered significant with dPSI>=0.1 and probability>=0.9. Gene ontology (GO) term enrichment was performed using the R package clusterProfiler (Yu et al. 2012).

### Quantitative RT-PCR

For HeLa and K562 cells total RNA was isolated using RNeasy Mini kit (Qiagen (Cat 74106)) and reverse transcribed with Transcriptor Universal cDNA Master (Roche). qRT-PCRs were run with standard settings on the Lightcyler 480 (Roche). Primers were designed using Roche Real-Time Ready Configurator combined with Roche Universal Probe Library (See Supplemental Table 4). Gene expression data was analysed by the delta Ct method normalised to the housekeeping gene POL2RJ, ACTB and MRIP. RT-PCR to validate splicing changes was performed using GoTaq (Promega) and quantified with Bioanalyzer RNA 6000 Nano assay (Agilent).

### CD34^+^ isolation from umbilical cord blood cells (UCB)

Cord blood samples were purchased from Anthony Nolan. Mononuclear cells (MNCs) were isolated from cord blood cells by centrifugation using Ficoll-PaqueTM PLUS (GE Healthcare Life Sciences, Buckinghamshire, UK). CD34^+^ cell enrichment was performed using EasySep™ Human CD34 Positive Selection Kit II (StemCell Technologies, Cat 17856) according to the manufacturer’s instructions.

### Lentivirus production in HEK 293T

Two lentiviral vectors, shRNA_DHX34#1 (GGAGCACGGATTGTGAATAAA), shRNA_DHX34#2 (GCCGACCAGGACAAGGTATTT) targeting human *DHX34* gene, and one control Scramble sequence (shRNA_Control) (CCTAAGGTTAAGTCGCCCTCG), were purchased from Vectorbuilder. All vectors were expressing GFP sequence as reporter gene. Viral particles for all the shRNAs were produced by transient CaCl2 transfection of HEK293T cells and harvested by ultracentrifugation.

### CD34^+^ UCB cells transduction

Umbilical cord blood (UBC) CD34^+^ HSPCs were stimulated using StemSpan medium (Stem cell Technologies, Cat 09655) supplemented with cytokines (150 ng/ml SCF, PeproTech, Cat 300-07; 150 ng/ml Flt-3, PeproTech, Cat 300-19; 10 ng/ml IL-6, PeproTech, Cat 200-06; 25 ng/ml G-CSF, PeproTech Cat 300-23; 20 ng/ml TPO, PeproTech Cat AF-300-18) and 1% HEPES (Sigma-Aldrich, Cat H0887-100mL) for 4-6 hours. Virus particles were then added to the stimulated cells (Multiplicity of infection, MOI=30) and cells were incubated (37ºC) overnight. Cells were washed and resuspended in expansion medium i.e (Stem cell Technologies, Cat 09655) with cytokines (150 ng/ml SCF, PeproTech, Cat 300-07; 150 ng/ml Flt-3, 20 ng/ml TPO, PeproTech Cat AF-300-18) and 1% HEPES (Sigma-Aldrich, Cat H0887-100mL). Cells were expanded for 4 days. Following on, cells were stained with antibody specific for human antigen CD34, DAPI (4,6, diamidino-2-phenylindole, Sigma-Aldrich, Cat D9542) staining was used to exclude dead cells and debris from the analysis. CD34^+^GFP^+^ cells were FACS sorted and then used in the different assays.

### RT-qPCR in HSPCs

RNA was extracted using RNeasy Mini Kit from Qiagen (Cat 74106) and retro-transcribed with High-Capacity cDNA Reverse Transcription Kit (ThermoFisher, Cat 4368814). qPCR was performed with Taqman probe for DHX34 (Hs00991248_m1, cat # 4351372, ThermoFisher) using B2M as endogenous control (Hs00984230_m1, cat#4331182, ThermoFisher).

### Colony forming assay

Two hundred and fifty patient CD34^+^ HSPCs were seeded in 0.5 mL methocult H4434 (StemCell Technologies, Cat 04434) supplemented with 1% penicillin/streptomycin (Sigma-Aldrich, Cat P4333) in a 24-well plate. Colonies were grown under hypoxic conditions (37ºC and 3% O_2_). Following 14 days of culture, colonies were scored.

### Erythroid differentiation

Transduced and FACs sorted CD34^+^ HSPCs were cultured in erythroid differentiation medium (SCF 25ng/mL, PeproTech, Cat 300-07; EPO 3U/mL, PeproTech, Cat 100-64; IGF1 50ng/mL, PeproTech, Cat 100-11) for 14 days. Cells were stained with antibodies specific for human antigens (CD71 PE RRID:AB 2201481; CD235a APC/Cyanine7, RRID:AB_ 2650977) and DAPI. Cells were immunophenotyped by using Fortessa flow cytometer (BD Biosciences, Oxford, UK) at day 4, 7, 10 and 14.

### Granulocytic differentiation

CD34^+^GFP^+^ HSPCs from UCB were cultured in granulocytic differentiation medium (SCF 25ng/mL, PeproTech, Cat 300-07; GM-CSF 10ng/mL, PeproTech, Cat 300-03) for 14 days. Cells were stained with antibodies specific for human antigens (CD11b APC RRID:AB_10561676; CD14 PE-Cy7, RRID:AB_1582277; CD45 APC eFluor780, RRID:AB_ 1944368) and DAPI. Cells were immunophenotyped by using Fortessa flow cytometer (BD Biosciences, Oxford, UK) at day 14.

### Cell cycle and Apoptosis

Cell were fixed/permeabilized with BD Cytofix/Cytoperm™ Kit (Cat 554714) and stained with DAPI (4,6, diamidino-2-phenylindole, Sigma-Aldrich, Cat D9542). DAPI (1 in 100 dilution) was used to assess cell cycle upon expansion conditions at day 14. Alexa Fluor® 647 Annexin V (Biolegend, Cat 640912) was used with Annexin V Binding Buffer (BD Bioscience, Cat 556454) to measure apoptosis at day 3 and 14 in expansion medium. Cells were analysed on Fortessa flow cytometer (BD Biosciences, Oxford, UK).

### Cell expansion

Cells were cultured in expansion media and cell number was measured at day 7 and 14 using a Countess 3 Automated Cell Counter.

### Statistical analysis

Prism Version 8 software (GraphPad) was used for statistical analysis in Figures 1, 3 and 6. Data are displayed as the mean±s.e.m. Statistical analysis was performed using unpaired two-tailed t-test for comparison of two groups. RStudio was used for the statistical analysis in Figures1, 2, 3 and 4 and Supplemental Fig. 1, 2, 3, 4 and 6. For information about the number of replicates, see the corresponding figure legend. For information about how data was analyzed and/or quantified, see the relevant section in Materials and methods.

## DATA DEPOSITION

All RNA-Seq data has been deposited in the Gene Expression Omnibus (GEO) database and an accession number will be available shortly.

## ACKNOWLEDGMENTS

We are grateful to Laura Monaghan (MRC HGU) for discussions and critical reading of the manuscript. We thank Sandra Keiper and Reinhard Luhrmann for discussions about RNA helicases and the spliceosome. M.R. was supported by funds from the Erasmus program. This work was supported by Core funding to the MRC Human Genetics Unit from the Medical Research Council (J.F.C) and by Blood Cancer UK (14032) abd Cancer Research UK (C15966/A24375) (J.F.; H.A.; A.R-M.) and by Kay Kendall Leukemia Fund (KKLF1149; K.R-P.).

## Author contributions

N.H. and J.F.C. conceived and designed the project. N.H performed the majority of experimental work, including CLIP experiments, DGE and AS analysis. M.R. contributed to the generation of tagged cell lines, interactome and CLIP experiments. A.R.M. performed the analysis of DHX34 pre-mRNA splicing. S.A. carried out the bioinformatic analysis. D.L performed data curation and analysis and participated in the draft of the manuscript and discussions. H.A, A.R.M., K.R-P. and J.F. carried out the biological experiments assessing the role of DHX34 in differentiation of erythroid and myeloid lineages. The manuscript was co-written by all authors.

## SUPPLEMENTAL MATERIAL

Supplemental material is available for this article.

