## Supplementary material for "A dual role for the RNA helicase DHX34 in NMD and pre-mRNA splicing and its function in hematopoietic differentiation": Supplememtal material

Running title: DHX34 role in pre-mRNA splicing and its impact in AML/MDS

**Supplemental Material includes:**

Supplemental Figures S1-S6

Supplemental Tables 1-4

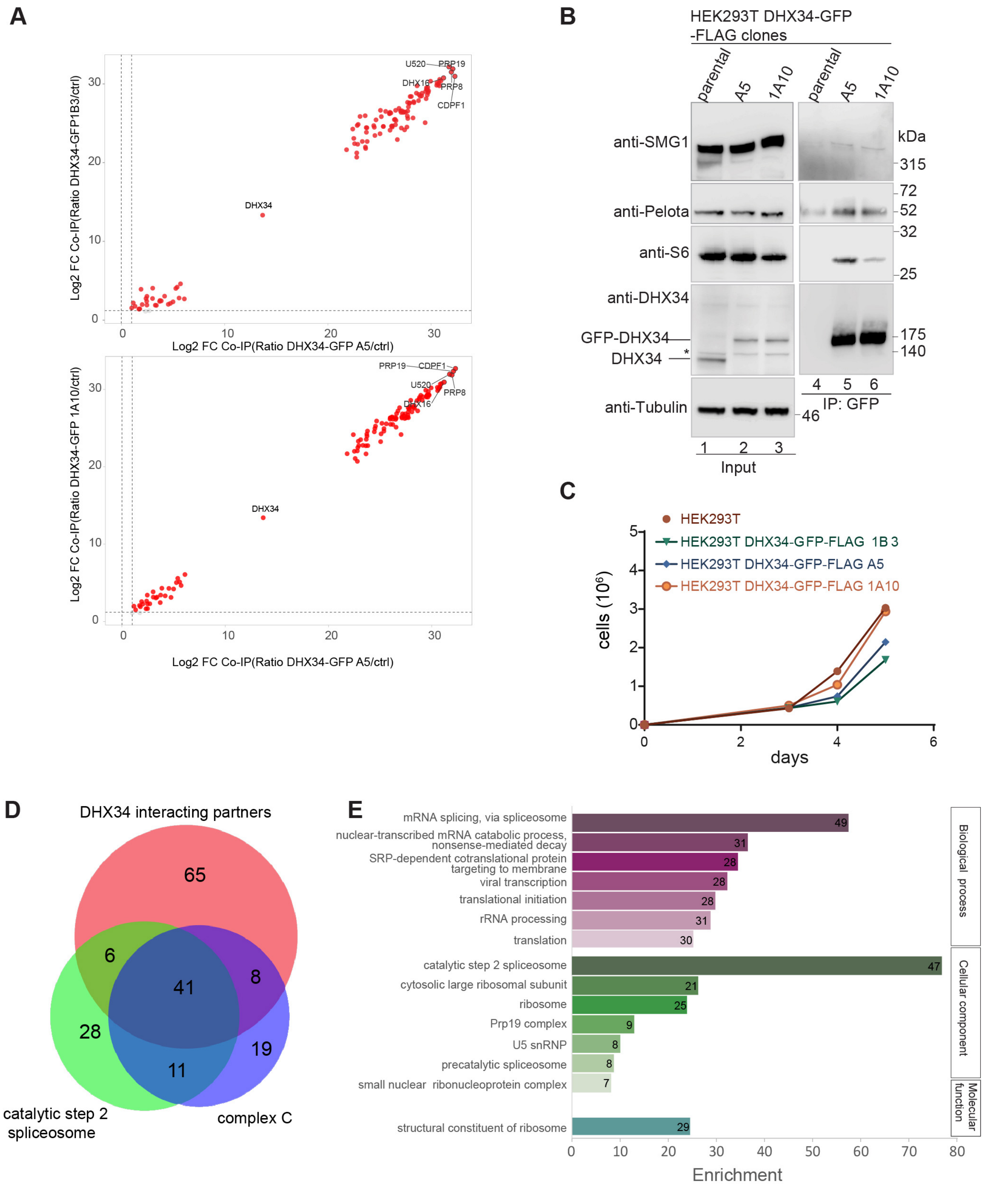


**Supplemental Figure S1.** DHX34 interacts with NMD components and mRNA processing complexes. Related to Figure 1. (*A*) Scatterplot showing overlap of protein enrichment ratios for DHX34-FLAG-GFP clones A5/1A10 and A5/1B3. (*B*) Validation of mass spectrometry results using IP (Immunoprecipitation)-Western blot analysis. Inputs and anti-GFP IP of parental cells and CRISPR clones used for mass spectrometry were separated by SDS-PAGE and probed with the indicated antibodies (*C*) Cell growth assay showing exponential growth for DHX34-FLAG-GFP and parental HEK293T cell lines. (*D*) DHX34 interacts with components of the spliceosomal C complex. Reactome and pathway analysis shows that DHX34 interacting partners are enriched for factors required for the second catalytic step of the splicing reaction and overlap largely with factors identified as a part of the late spliceosomal complex C. (*E*) GO term enrichment of DHX34 interacting partners.

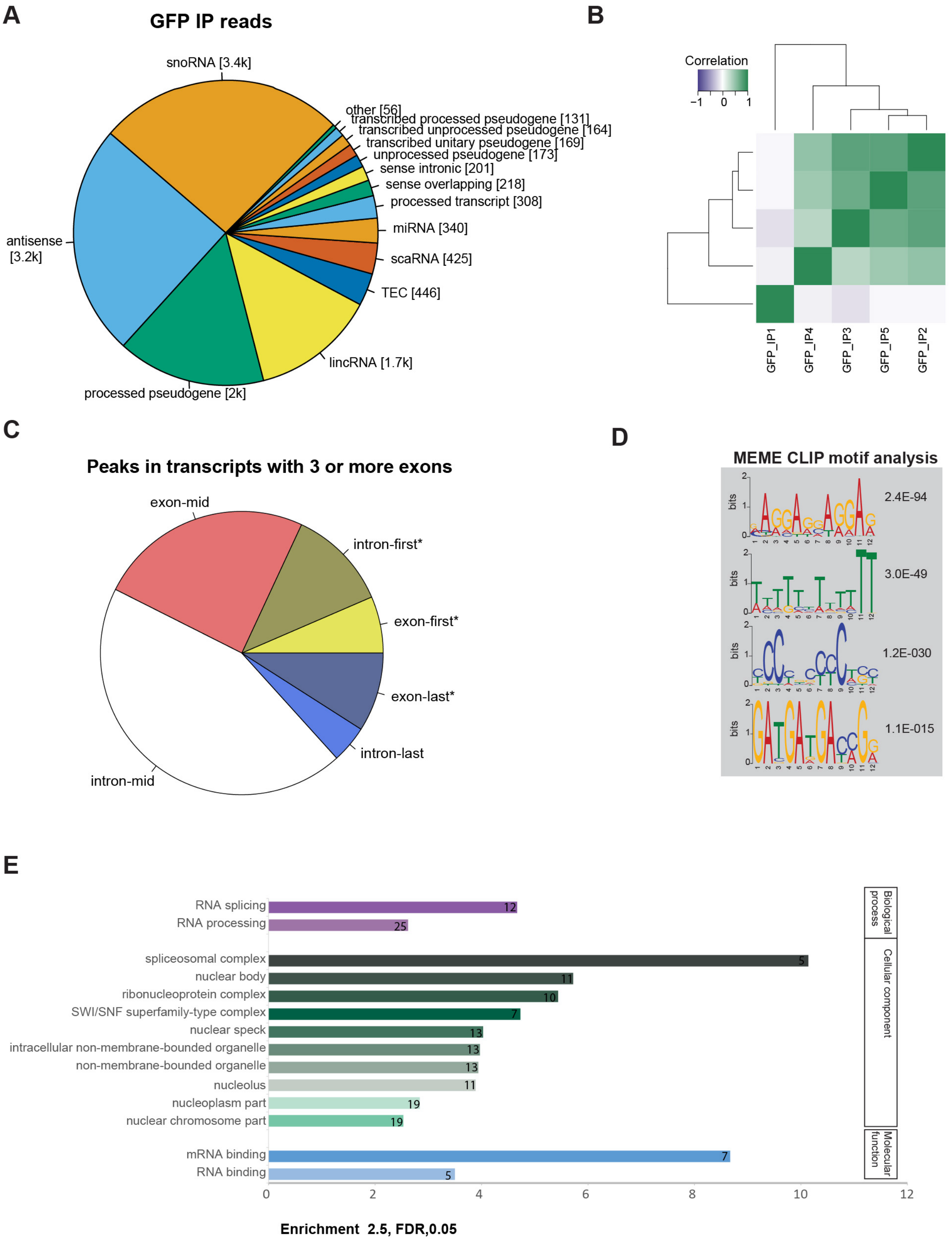


**Supplemental Figure S2.** RNA-binding pattern of DHX34. Related to Figure 2. (*A*) Pie chart showing the distribution of non-protein coding transcripts in DHX34 RNA targets. Total normalized reads mapping to gene bodies per biotype is shown in pie_chart_biotypes_GFP_IP.pdf for all GFP IP reads, and when summing reads over those genes where GFP IP is 1.5 times enriched over negative GFP IP and over the input. (*B*) Correlation of raw counts for the selected peaks across replicates. Heatmap of selected peaks which were supported by 5-11 raw reads across replicates raw counts. Peaks in GFP_IP1 though strongly indicated, correlated less well. (*C*) Metagene analysis investigating the position of DHX34 RNA-binding in protein coding transcript expressing more than three exons. Higher-than-expected binding frequency is indicated by asterisk (*D*) MEME motif analysis of seCLIP peaks. The four most significant motifs for DHX34 are shown. (*E*) GO term enrichment (enrichment ≥2.5, FDR≤0.05) for genes with DHX34 RNA-binding peaks in seCLIP.

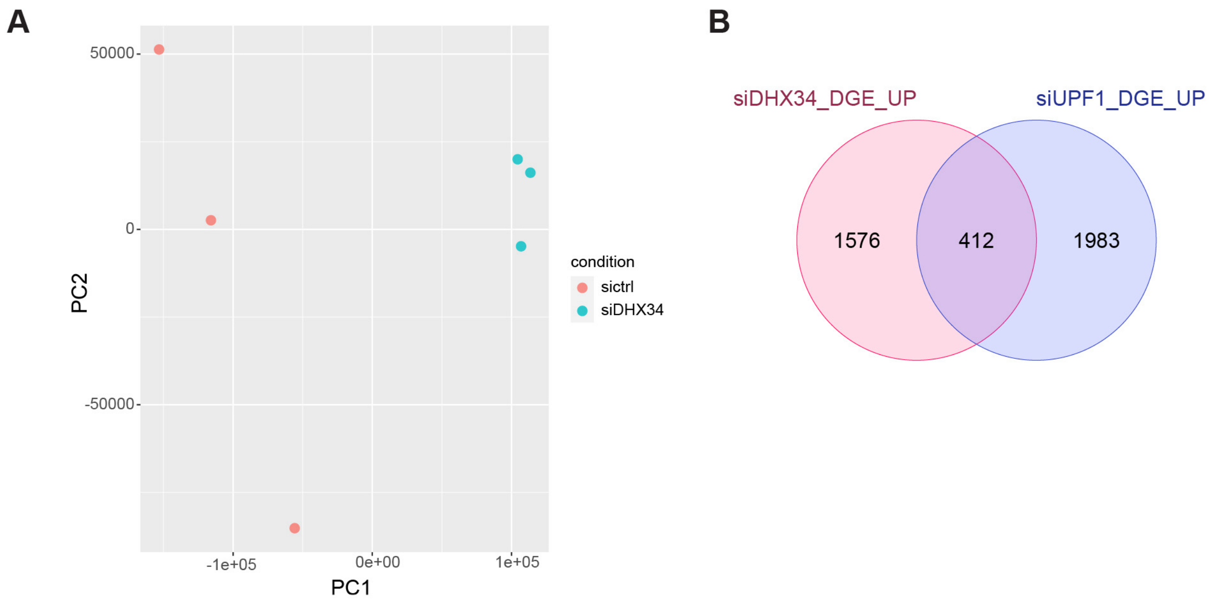


**Supplemental Figure S3.** DHX34 affects mRNA expression. Related to Figure 3. (*A*) Principal component analysis (PCA) of differential gene expression data. (*B*) Overlap of genes upregulated upon DHX34 depletion (this study) compared to UPF1 knockdown (Longman et al. 2020) is shown.

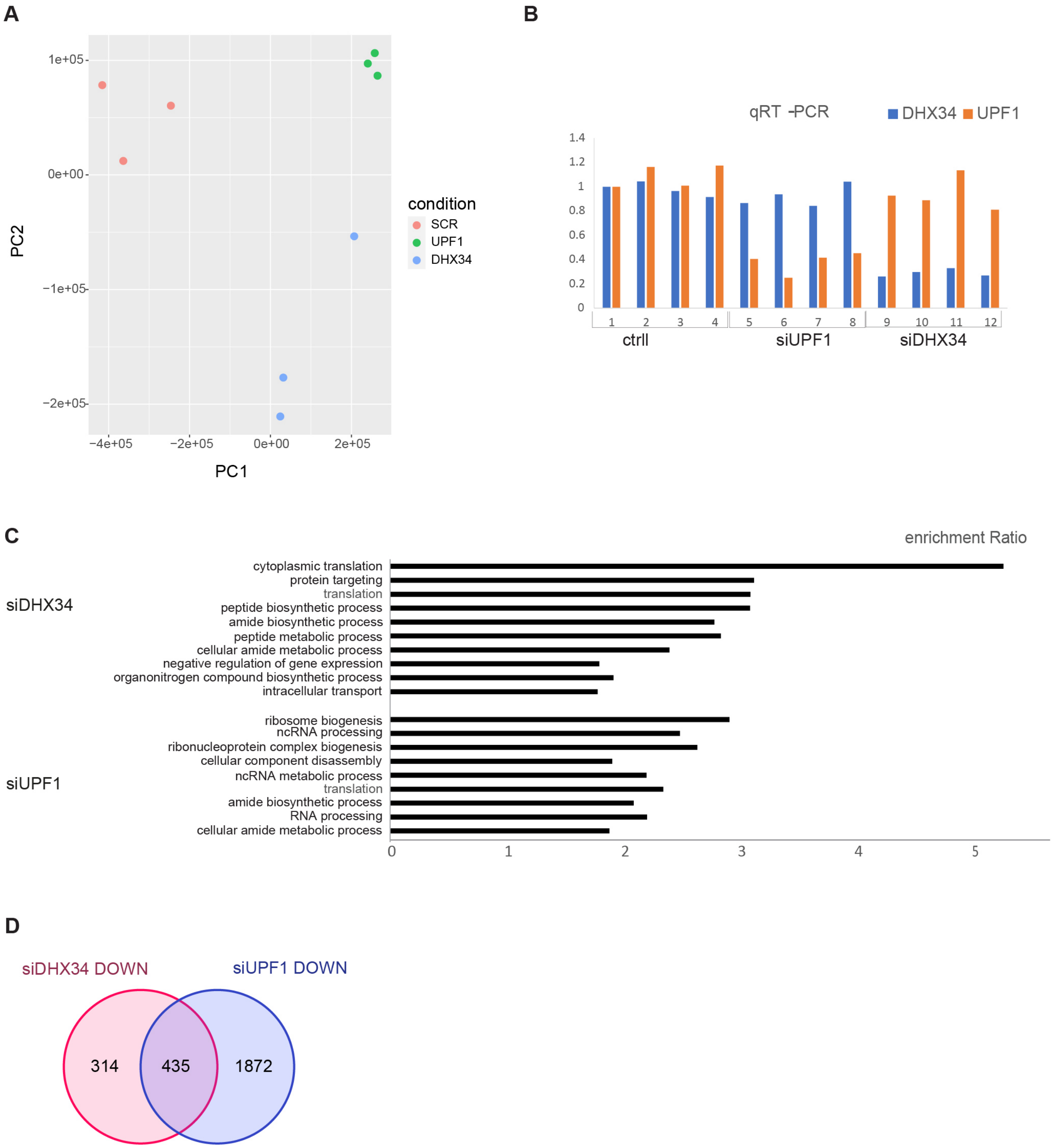


**Supplemental Figure S4.** DHX34 affects gene expression in K562 cells. Related to Figure 4. (*A*) Principal component analysis (PCA) of differential gene expression data for knockdown experiments in K562 cells. (*B*) qRT-PCR validation of DHX34 and UPF1 knockdown compared to non-targeting siRNA (ctrl). In total, 3 out of 4 biological replicas were used for RNA seq. (*C*) GO term enrichment (enrichment ≥2.5, FDR≤0.05) for genes upregulated upon DHX34 and UPF1 knockdown. (*D*) Venn diagram showing the number of common down-regulated targets (DOWN) in DHX34 and UPF1 knockdown cells.

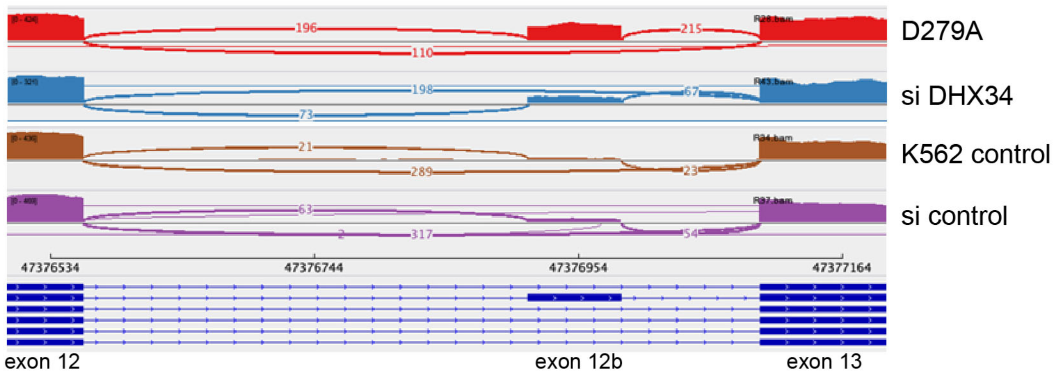


**Supplemental Figure S5.** DHX34 affects alternative splicing in K562 cells. Related to Figure 5. Sashimi plot of control depleted and mutant samples visualizing splicing junctions of DHX34 between exons 12 and 13 from aligned RNA seq data.

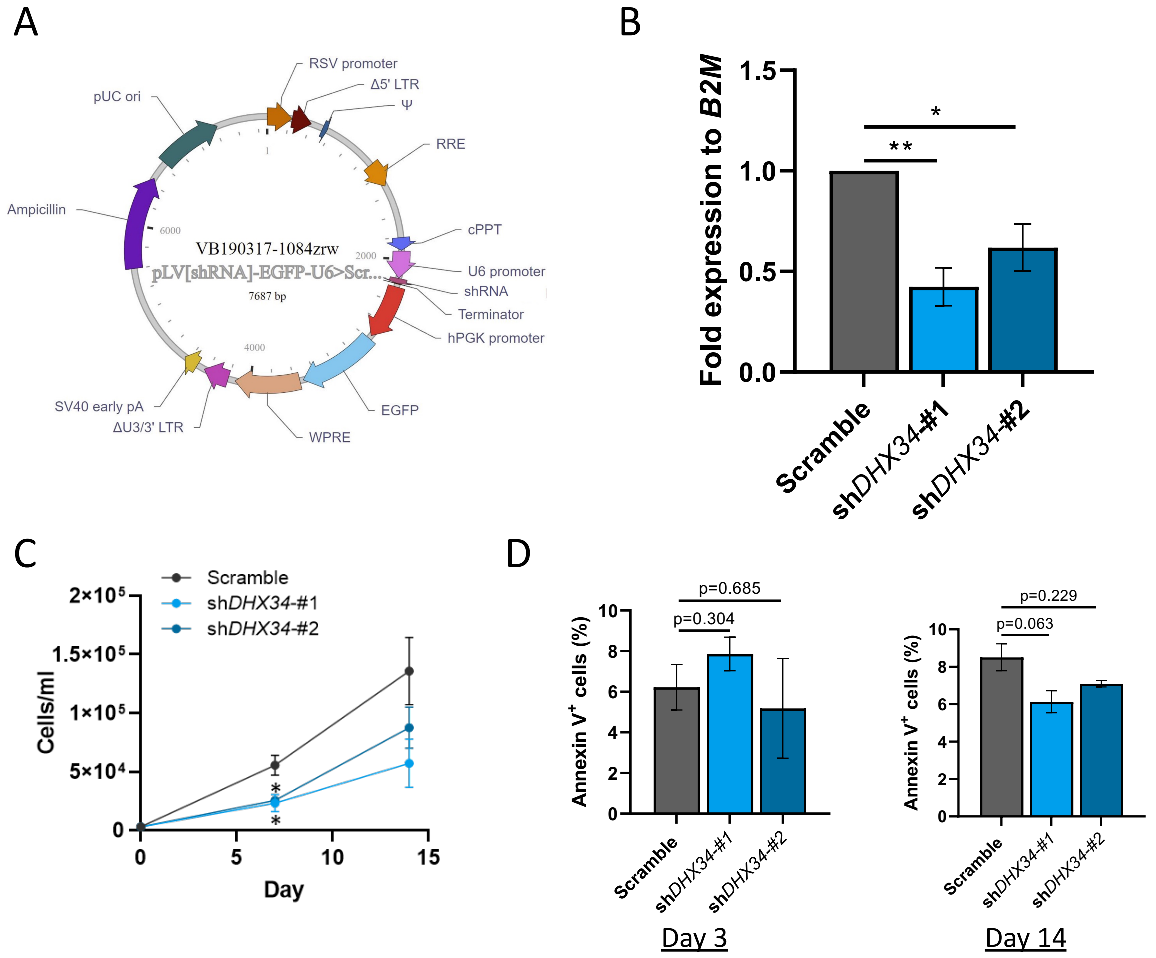


**Supplemental Figure S6.** Loss of DHX34 impairs HSPCs differentiations. Related to Fig. 6. *(A)* Schematic representation of the lentiviral plasmid used to clone the shRNA, which is under the control of U6 promoter, whereas the eGFP reporter gene is under the control of the PGK promoter. (*B*) RT-qPCR of *DHX34* upon KD with two different shRNAs in primary CD34 cells 4-days post-transduction. Expression is normalized to *B2M* (n=3). (*C*) Cells were sorted by flow cytometer (DAPI$^-$CD34$^+$GFP$^+$) and placed in expansion medium to be scored at day 7 & 14 (n=3). (*D*) Apoptosis was assessed in cells cultured in expansion medium by FACS immunophenotyping with Annexin V antibody at day 3 and day 14 (n=3). *=p<0.05; **=p<0.01; ***=p<0.005

**Supplemental Table 1.** List of proteins identified by Mass-spec following anti-GFP immunopurifications of C-terminal tagged DHX34-GFP-FLAG cell lines. Related to Figure 1.

**Supplemental Table 2.** Changes in DGE and AS in HeLa cells upon DHX34 KD. Related to Figure 3.

**Supplemental Table 3.** Changes in DGE and AS in K562 cells upon DHX34 KD and DGE upon UPF1 KD. Related to Figure 4.

**Supplemental Table 4.** List of siRNAs and RT-PCR primers.
