## Supplemental Table 4 for "A dual role for the RNA helicase DHX34 in NMD and pre-mRNA splicing and its function in hematopoietic differentiation"

Supplemental Table 1

primers

| use | name | sequence |
| --- | --- | --- |
| RT-PCR | DHX34 fwd | TTTCAGCAATGCCCCTGT |
| RT-PCR | DHX34 rev | CTCCTGCGGCTGGTACAC |
| RT-PCR | UPF1 fwd | CCTGCACACCAAGCTCTACC |
| RT-PCR | UPF1 rev | TTCTCGCGTCCCTGAAAG |
| RT-PCR | ACTB fwd | AGAGCTACGAGCTGCCTGAC |
| RT-PCR | ACTB rev | CGTGGATGCCACAGGACT |
| RT-PCR | MRIP fwd | CGAAGGCTAAGGCTGACTGT |
| RT-PCR | MRIP rev | ACTTCTCCCCCAGTGCTTC |
| RT-PCR | POLR2J fwd | CTGTGAGCCCCGTTCCTAC |
| RT-PCR | POLR2J rev | GTCGGTGTCAGGGTGAGG |
| RT-PCR | DHX34 ex12 F | TTT CCA CAC GCA GGC CAA GCA GGG CG |
| RT-PCR | DHX34 ex12 R | CAG GGA TGC GGA CGC AGT TCA CCA GG |
| RT_PCR | U1_L1 | AAGGTGGTTTTCCCAGGGCG |
| RT_PCR | U1_R1 | CGAACGCAGTCCCCCACTAC |
| RT_PCR | U2_L1 | TCGCTTCTCGGCCTTTTGGC |
| RT_PCR | U2_R1 | GATGCGTGGAGTGGACGGAG |
| RT_PCR | U4_L1 | AGCTTTGCGCAGTGGCAGTA |
| RT_PCR | U4_R1 | ATTGCAAGTCGTCACGGCGG |
| RT_PCR | U5_L1 | ACTCTGGTTTCTCTTCAGATCGCA |
| RT_PCR | U5_R1 | CTTGCCAAAGCAAGGCCTCA |
| RT_PCR | U6_L1 | GCTCGCTTCGGCAGCACATA |
| RT_PCR | U6_R1 | TATGGAACGCTTCACGAATTTGC |
| RT_PCR | ACOT9_F | CAAAGGGCAGCTTACTCCTGG |
| RT_PCR | ACOT9_R | TGCTCCTACTATCTCCCGCAA |
| RT_PCR | CLSTN1_F | TGTGACTGAGGATTACCCGCT |
| RT_PCR | CLSTN1_R | GAACGGAGAGTTAAGCCAGCC |
| RT_PCR | MAP2K7_F | AAAGCTGAAGCAGGAGAACCG |
| RT_PCR | MAP2K7_R | GAGCTCTCTGAGGATGGCGAG |
| RT_PCR | MYL6_F | AGAAGTAGAGATGCTGGTGGC |
| RT_PCR | MYL6_R | ATTCACACAGGGAAAGGCACG |
| RT_PCR | MYOF_F | AAAATTGCTGCCTCTGGTGGG |
| RT_PCR | MYOF_R | AATCCCGTGTACTCTCTGGGG |

| guide RNA | Sequence | backbone | Reference |
| --- | --- | --- | --- |
| DHX34 C-term | GGAAGCAGCACGTGTGAGCTGGG | pSpCas9(BB)-2A-Puro (px459) V2.0 | pSpCas9(BB)-2A-Puro (PX459) V2.0 was a gift from Feng Zhang (Addgene plasmid # 62988 ; http://n2t.net/addgene:62988 ; RRID:Addgene_62988) |
| D279A | CCAGTATGAGGTCCTGATTGTGG | pSpCas9(BB)-2A-Puro (px459) V2.0 |  |
| E444D | GACTGAGGTCTCAGCAATGTTGG | pSpCas9(BB)-2A-Puro (px459) V2.0 |  |
| Y515C | CCTACCCCGTCCCAGAAATTCGG | pCMV_AncBE4max_P2A_GFP | was a gift from David Liu (Addgene plasmid # 112100 ;http://n2t.net/addgene:112100 ; RRID:Addgene_112100) |

| **ss oligo HR template** |  |  |
| --- | --- | --- |
| D279A_HR | sequence can be given upon request |  |
| E444D_HR | sequence can be given upon request |  |
| **siRNA** |  |  |
| scramble | siGENOME No-Targeting siRNA Pool 1 | D-001206-13-20 (Dharmacon) |
| DHX34 pool | siGENOME Human DHX34 siRNA | M-032233-01-0010 (Dharmacon) |
| UPF1 siRNA I, II |  | Described in Longman et al. 2020 |
